# Towards On-Site Paleogenomics: Application and Perspective of Nanopore Sequencing with Ancient DNA

**DOI:** 10.64898/2025.12.29.696801

**Authors:** Koji Ishiya, Ravdandorj Odongoo, Kenji Kasai, Kenichi Machida, Shigeki Nakagome, Shoko Tarumoto, Takuya Yamamoto, Taro Tsujimura, Takashi Gakuhari

**Affiliations:** Sapiens Life Sciences, Evolution and Medicine Research Center, Kanazawa University, Ishikawa, JAPAN; Institute for the Study of Ancient Civilizations and Cultural Resources, Kanazawa University, Ishikawa, JAPAN; Toyama Prefectural Center for Archaeological Operations, Toyama, JAPAN; Toyama Prefectural Research Office for Archaeological Heritage, Toyama, JAPAN; School of Medicine, Trinity College Dublin, Dublin, IRELAND; Institute for the Advanced Study of Human Biology (WPI-ASHBi), Kyoto University, Kyoto, JAPAN; Center for iPS Cell Research and Application (CiRA), Kyoto University, Kyoto, JAPAN

**Keywords:** Ancient DNA, paleogenomics, nanopore sequencing, ethical DNA research

## Abstract

Ancient DNA (aDNA) research has greatly advanced understanding of past populations, yet progress in this field is still limited by two fundamental issues: dependence on fixed laboratory infrastructure and ethical considerations surrounding the cross-border transfer of archaeological biological specimens. In this study, we present the first successful application of Oxford Nanopore Technologies (ONT) sequencing to authentic aDNA from ancient human remains dating to the Early Jomon period. Our results demonstrate that nanopore platforms can recover characteristic postmortem damage signatures, generate genome-wide information with time-stamped sequencing data, and produce population genetic inferences consistent with Illumina short-read data. Crucially, the ONT sequencing has enabled the first demonstration of time-resolved aDNA analysis. This enables key genetic metrics, such as biological sex inference, to be determined within the first 60 minutes of sequencing. The portability and operational simplicity of ONT devices provide a practical basis for on-site paleogenomics, facilitating the generation of genomic data directly at archaeological sites, in museums, or at local research institutions. This capability is particularly significant in regions where the export of biological specimens entails administrative procedures, often spanning several months or even years. By enabling local archaeologists and anthropologists to independently generate and interpret genomic data, on-site nanopore sequencing can accelerate global research alliance, and promote equitable scientific authorship and continuing dialogue on sample sovereignty and collaborative research practices between geneticists and field researchers. Together, our findings establish nanopore sequencing as a socially sustainable and viable tool for paleogenomics, offering new pathways for the rapid, field-deployable and ethical process of archaeological remains.

## 1. Introduction

Ancient DNA (aDNA) research has profoundly transformed archaeology, anthropology, and evolutionary biology since its inception nearly four decades ago [1–3]. aDNA provides a direct means to reconstruct evolutionary history, functional adaptations, and population dynamics of past organisms, including archaic hominins [4–7], humans [8–11], domestic animals [12–14], plants [15–17], microbes [18–22], and viruses [23]. Advances in next-generation sequencing (NGS) technologies [24] have dramatically increased the scale and resolution of paleogenomic studies, enabling researchers to recover detailed genetic information from increasingly older and more degraded specimens.

Despite these advances, most aDNA studies continue to rely on stationary short-read sequencing platforms such as Illumina’s sequencing-by-synthesis (SBS) system [25]. These platforms deliver high throughput and accuracy but require dedicated laboratory infrastructure and cannot be deployed in the field. In addition, international regulations governing the movement of archaeological remains [26]—including extended approval processes and export restrictions—can pose logistical challenges for cross-border collaborations. These factors may limit opportunities for local archaeologists and anthropologists to play a leading role in genomic studies of their own cultural heritage.

Oxford Nanopore Technologies (ONT) [27] offers an alternative model through highly portable, real-time sequencing systems capable of generating long single-molecule reads. ONT instruments have transformed genome assembly [29–31], transcriptomics [32, 33], epigenomics [34], and metagenomics [35–37], yet their applicability to paleogenomics has remained unclear. aDNA molecules are typically short, chemically damaged, and structurally unstable, raising concerns that ONT’s sequencing chemistry and basecalling algorithms may be incompatible with authentic aDNA signatures. If validated, ONT sequencing could enable on-site paleogenomics, allowing local researchers to generate genomic data without exporting archeological materials. This shift would accelerate data accessibility, reduce global inequities in scientific authorship, and alleviate long-standing tensions between archaeologists and geneticists regarding sample sovereignty, research leadership, and the stewardship of archelogical remains.

In this study, we present the first successful application of ONT sequencing to authentic ancient human DNA, demonstrating that nanopore platforms preserve canonical postmortem damage patterns and yield population-genetic results concordant with Illumina datasets. We further show that ONT’s time-resolved output enables rapid biological inference—such as sex determination—within the first hour of sequencing. These findings establish a foundation for flexible, field-deployable, and ethically sustainable paleogenomics.

## 2. Results

### 2.1 Sequencing summary

We generated whole-genome shotgun sequencing data for individual ODK14 using ONT PromethION 2 Solo (P2Solo) and Illumina Novaseq 6000. The P2Solo run produced 17.81 Gb of raw data, comprising 60,625,684 reads across 72 hours of sequencing (Fig. 1). Among these reads, 15,973,588 (26.35%) exhibited an average Phred score ≥Q20, and 11,933,911,534 bases (67.02%) exceeded Q20 at the per-base level (Fig. S1). Pore translocation profiling via Sequali indicated that 30.78% of reads fell within expected speed ranges, 67.00% were slower, and 2.22% were faster than expected (Fig. S2). The raw mean read length prior to QC filtering was 293.72 bp. Illumina Novaseq 6000 sequencing generated 10.1 Gb of paired-end reads (2 × 151 bp; 67,052,219 total reads). After adapter removal and trimming, QC-filtered ONT reads mapped to the human genome at 15.45%, while QC-filtered Illumina reads mapped at 18.23% (Table 1). We also detected 5-methylcytosine (5mC) at 3,101 CpG sites (0.19%) out of 1,593,565 CpG sites in the reference human genome (GRCh37) from the ONT sequencing data (Fig. S3). However, the majority of these sites had coverage of 5X or lower, which may lead to biased methylation rate estimates (Fig. S4);therefore, comparison with known methylated regions was considered inapplicable due to insufficient coverage.

**Figure 1.**
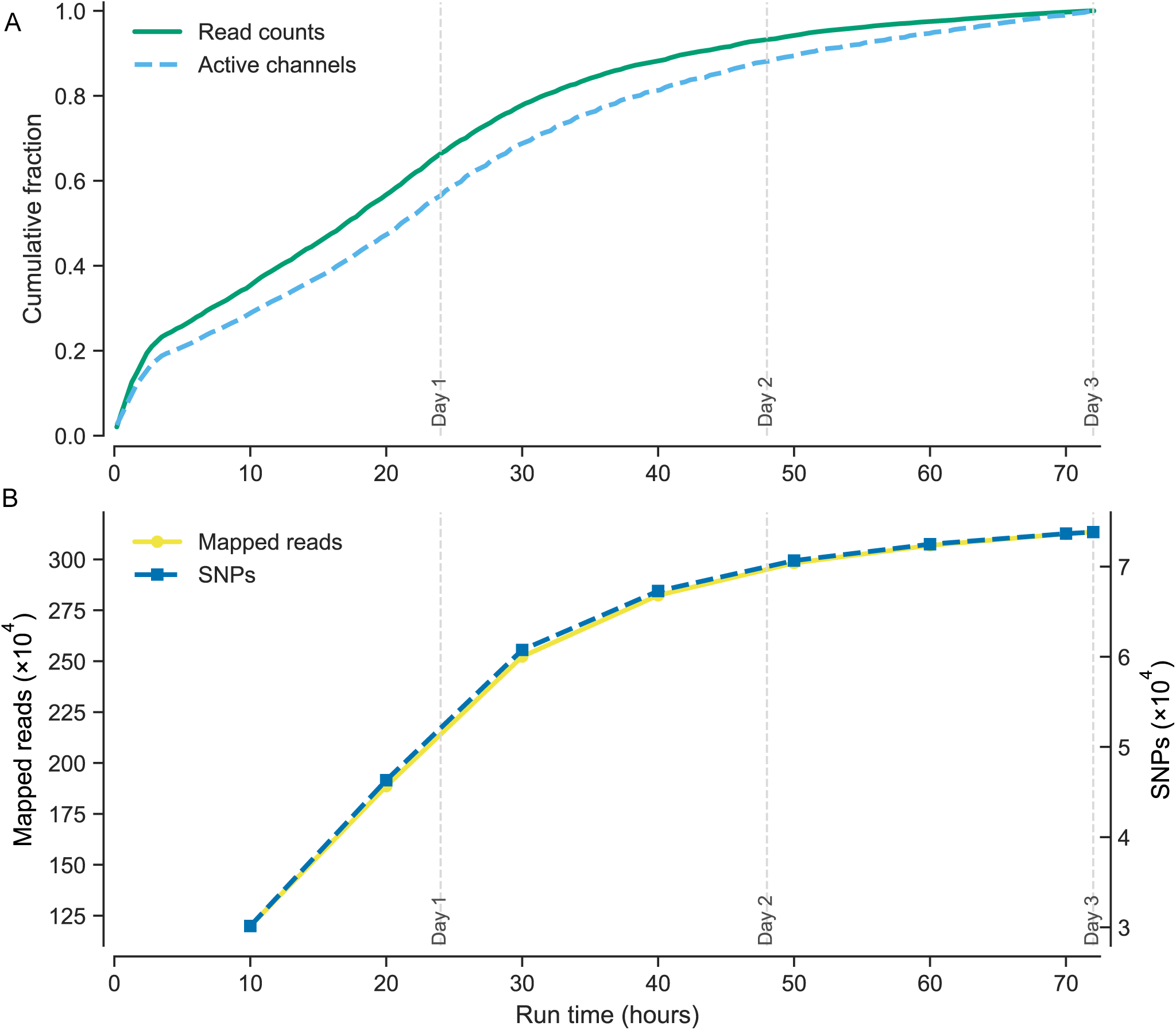
Temporal progression of sequencing output across ONT runtime intervals. This figure summarizes sequencing and genomic data generated from Oxford Nanopore Technologies (ONT) runs on an ancient DNA (aDNA) sample over successive intervals from 10 to 72 hours. (A) Cumulative plot showing the number of raw reads and active nanopore channels based on ONT sequencing statistics. (B) The yellow line represents reads successfully aligned to the human reference genome (hg19/GRCh37) with mapping quality ≥30 and duplicate removal, used for SNP genotype calling. The blue line indicates the cumulative number of observed SNPs within the 1240K panel.

**Table 1.**
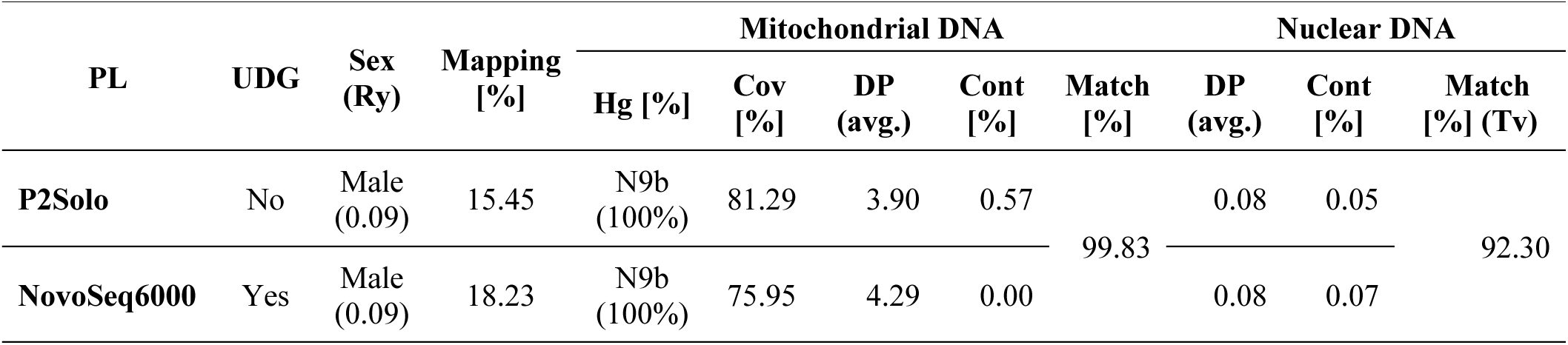
Sequencing summary from different platforms for ODK14. “PL” denotes the sequencing platform. “UDG” indicates whether uracil-DNA glycosylase treatment was applied (“No” indicates untreated libraries). “Hg” represents the mitochondrial haplogroup inferred from the consensus mitochondrial genome using MitoSuite, and the percentage indicates the concordance rate at haplogroup-defining sites. “Sex” indicates the biological sex estimated from the ratio of reads mapped to the X and Y chromosomes (Ry). “Cov” denotes the average coverage of the mitochondrial genome and autosomes (chr1–22), and “DP” refers to sequencing depth. “Cont” shows estimated contamination rates: mitochondrial contamination is based on haplogroup-defining sites, whereas nuclear contamination is estimated from X-chromosomal heterozygosity. “Match” indicates the concordance percentage between P2Solo and NovoSeq6000 sequencing results; for nuclear genome comparisons, this percentage was calculated using transversion (Tv) sites to minimize the impact of postmortem damage.

### 2.2 Ancient DNA authenticity in ONT reads

To confirm that the DNA recovered from the ONT sequencing reflects authentic ancient DNA (aDNA), we examined several characteristic damage profiles typically observed in ancient samples. Because the ONT reads were generated from a UDG-untreated extraction library, we observed pronounced deamination-derived substitutions accumulating toward both ends of the reads (Fig. 2A). The aligned ONT reads also showed substantial fragmentation, with an average read length of 79.5 bp (Fig. 2B). In addition, accumulated damage increased the number of mismatches between the reads and the reference genome, resulting in elevated edit distances (Fig. 2C). We also assessed exgenonus DNA contamination in the ONT reads, with both mitochondrial and X-chromosomal estimates falling below 1% (Table 1). Combined with the characteristic damage profiles, this low level of contamination further supports the authenticity of aDNA and demonstrates that these signals can be reliably recovered even using ONT sequencing.

**Figure 2.**
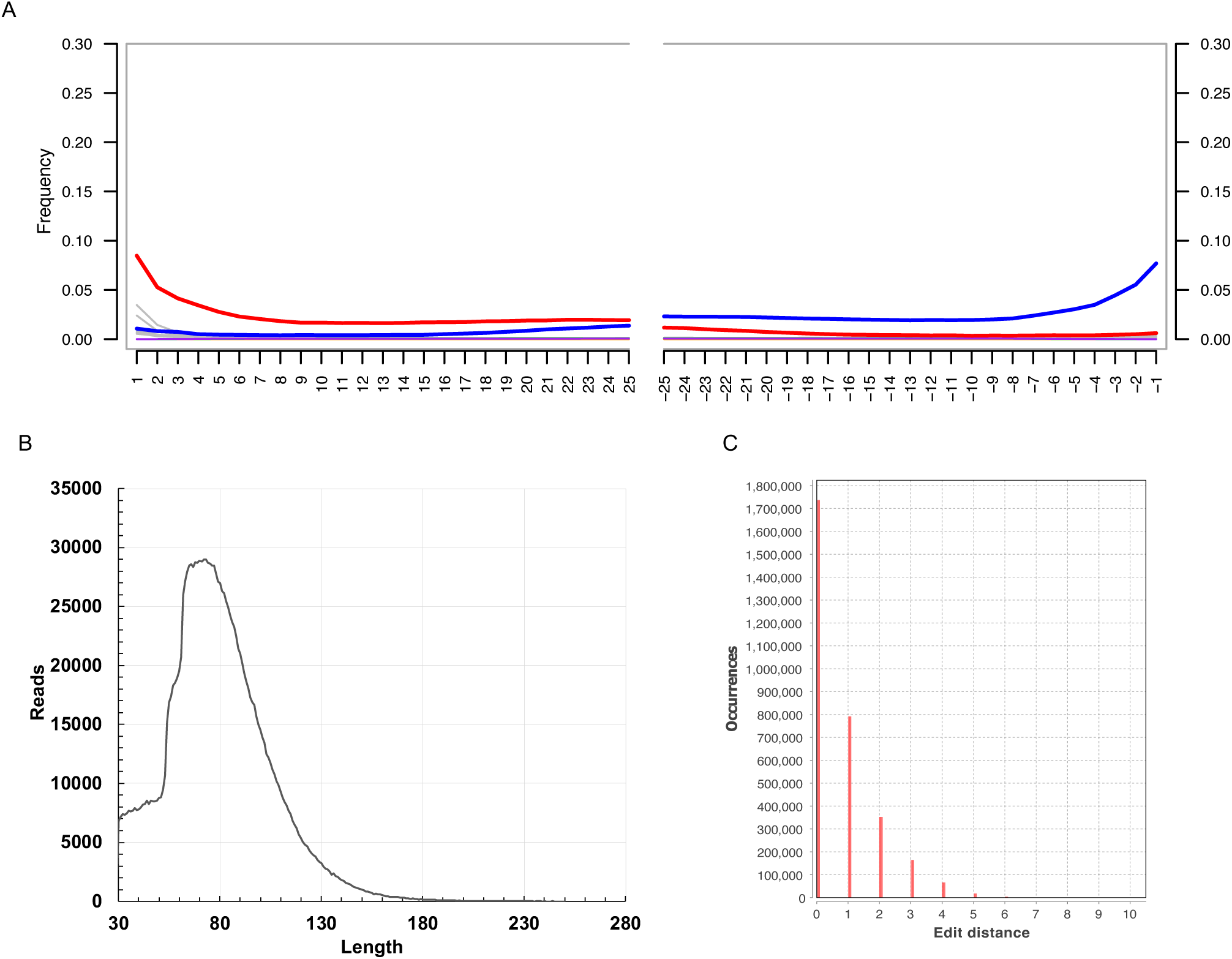
Characteristics of post-mortem damaged aDNA in ONT sequencing data. (A) Deamination patterns in mapped reads, as visualized by mapDamage. The x-axis indicates the distance from the read ends, while the y-axis shows the frequency of base substitutions. Red lines represent C→T substitutions at the 5’ end, blue lines represent G→A substitutions at the 3’ end, and other substitutions are shown in gray. (B) Distribution of DNA fragment lengths in mapped reads, illustrating DNA fragmentation profiles. (C) Edit distance between mapped reads and the reference human genome based on their observed mismatches, calculated using DamageProfiler.

### 2.3 Comparative evaluation with illumina sequencing data

To evaluate cross-platform concordance between Oxford Nanopore Technologies (ONT) and Illumina datasets, we examined the following: (i) maternal/paternal lineage assignment, (ii) genome-wide coverage, and (iii) single nucleotide polymorphism (SNP)-level agreement (Table 1). Since the ONT libraries were UDG-untreated to preserve cytosine deamination, the concordance analyses were limited to transversion (Tv) sites, which are minimally impacted by postmortem damage. Both platforms assigned ODK14 to the mitochondrial haplogroup N9b, which is frequently reported in Jomon individuals [38–42] (Table 1, Fig. S5). However, Y-chromosomal haplogroup inference was not feasible due to the insufficient recovery of informative SNPs. Mean mitochondrial coverage was slightly higher for Illumina (4.29×) than for ONT (3.90×); autosomal coverage was consistent across platforms at 0.08×. Across the transversion sites of the widely used 1240K SNP panel — comprising approximately 1.24 million SNPs for human population genetic analyses — ONT-based genotypes agreed with Illumina-based calls for 92.30% of sites (Table 1, Fig. S6). The residual discordance likely reflects differences in library construction and sequencing chemistry between the two platforms, as well as how they handle damage and basecalling. Overall, these results suggest that ONT-derived genotypes are generally consistent with Illumina-derived genotypes, supporting the reliability of subsequent paleogenomic inferences.

### 2.4 Time-resolved recovery of genomic data

Nanopore sequencing platforms such as MinION and P2Solo provide advantages not only in portability but also in their ability to record precise, time-resolved sequencing information [27] (Fig. 1). We leveraged this capability as a proof-of-concept to apply temporal analyses to aDNA sequencing. Using timestamps associated with each ONT read, we partitioned the dataset into discrete time intervals and evaluated how rapidly genomic data could be recovered, including metrics relevant to sex determination, alignment yield, and SNP detection.

Notably, the ratio of X- to Y-chromosomal reads (Ry) [43], used for biological sex inference in archaeological samples, converged to a stable estimate within the first 60 minutes of sequencing, supporting a male assignment for the ODK14 individual (Fig. 3). This estimate remained highly consistent throughout the entire sequencing run (∼72 hours; Table 1), indicating that reliable sex determination can be achieved from minimal ONT data even for degraded or archaeological samples.

**Figure 3.**
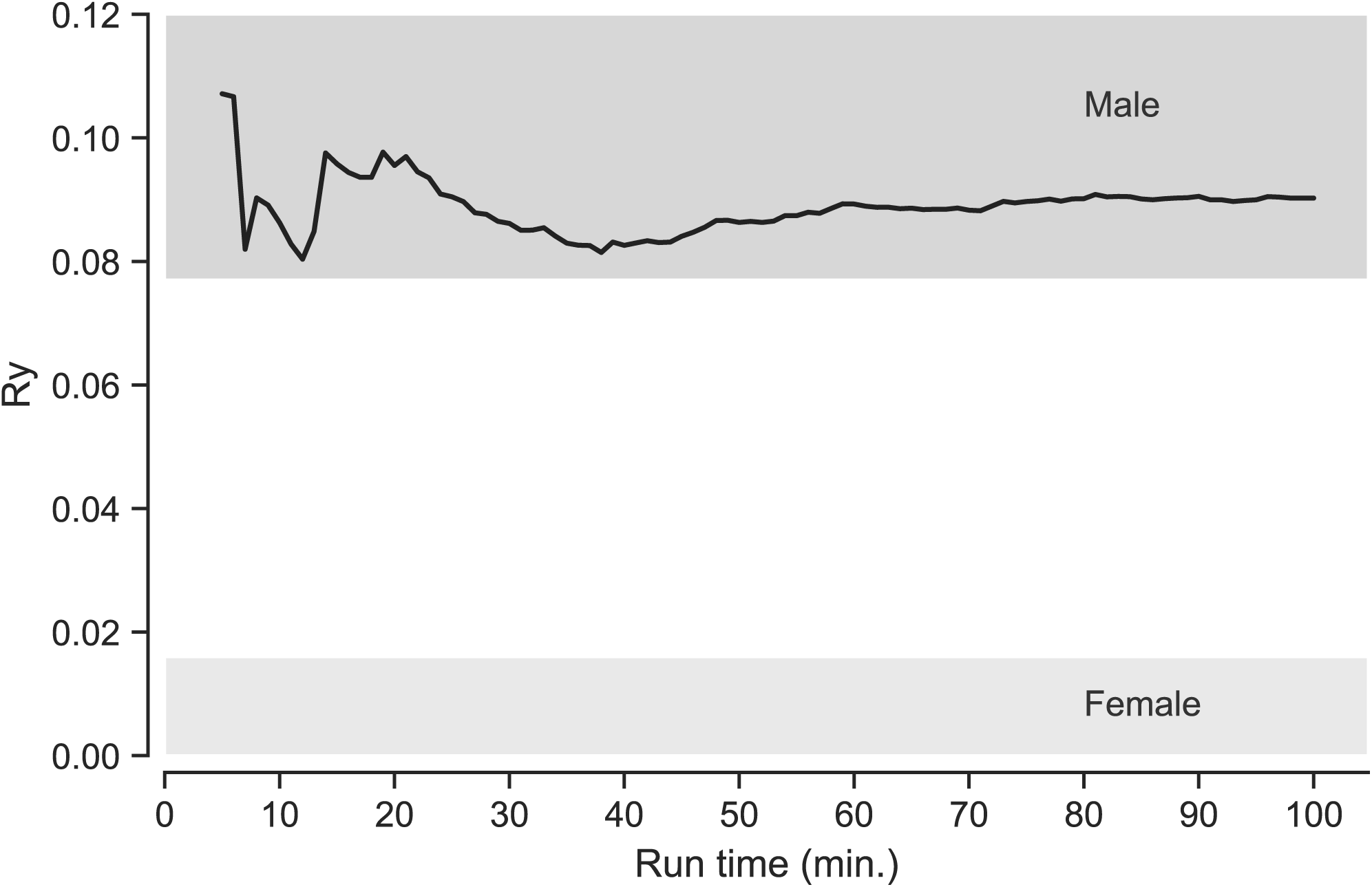
Temporal changes in Y/X+Y alignment ratio (Ry) during initial sequencing phase (0-100 minutes). Changes in Ry—the ratio of reads aligned to the Y chromosome versus the X chromosome—over the runtime (0–100 minutes). The shaded regions represent the male and female reference ranges for Ry values used for sex determination, following Skoglund et al. (2013).

The number of raw ONT reads increased rapidly during the first two days of sequencing, reaching nearly 90% of the total yield by the end of day two, with only a modest additional increase (∼10%) on the third day (Fig. 1A). This temporal pattern was mirrored in both the number of aligned reads and the number of recovered SNPs. In particular, the trajectories of aligned reads and SNP counts closely paralleled each other, increasing proportionally with the accumulation of raw reads up to the end of day two (Fig. 1B).

### 2.5 Population genetics analysis

To evaluate whether the ODK14 individual exhibits genetic affinities comparable to previously reported Jomon individuals sequenced using Illumina technologies, we conducted a series of population genetic analyses.

First, we assessed the genetic affinities of ODK14 among ancient and present-day populations across East Eurasia using outgroup *f3* statistics [44]. The results showed that ODK14 shares the highest affinity with previously published Jomon individuals, and among present-day East Eurasian populations, the strongest affinity was observed with modern Japanese (Fig. 4, Table S3). These findings indicate that ODK14 possesses genetic characteristics consistent with those of the Jomon lineage. We next performed principal component analysis (PCA) [45] using a panel dataset comprising ancient individuals and present-day East Eurasian populations. On the second principal component, published Jomon individuals occupy a distinct position separated from other East Eurasian groups, and ODK14 located within this same region (Fig. 5), further supporting its Jomon-related ancestry. ADMIXTURE analysis [46] identified *K* = 3 as the model with the lowest cross-validation error. Under the model, ODK14 shared similar ancestry components in previously reported Jomon individuals (Fig. 6).

**Figure 4.**
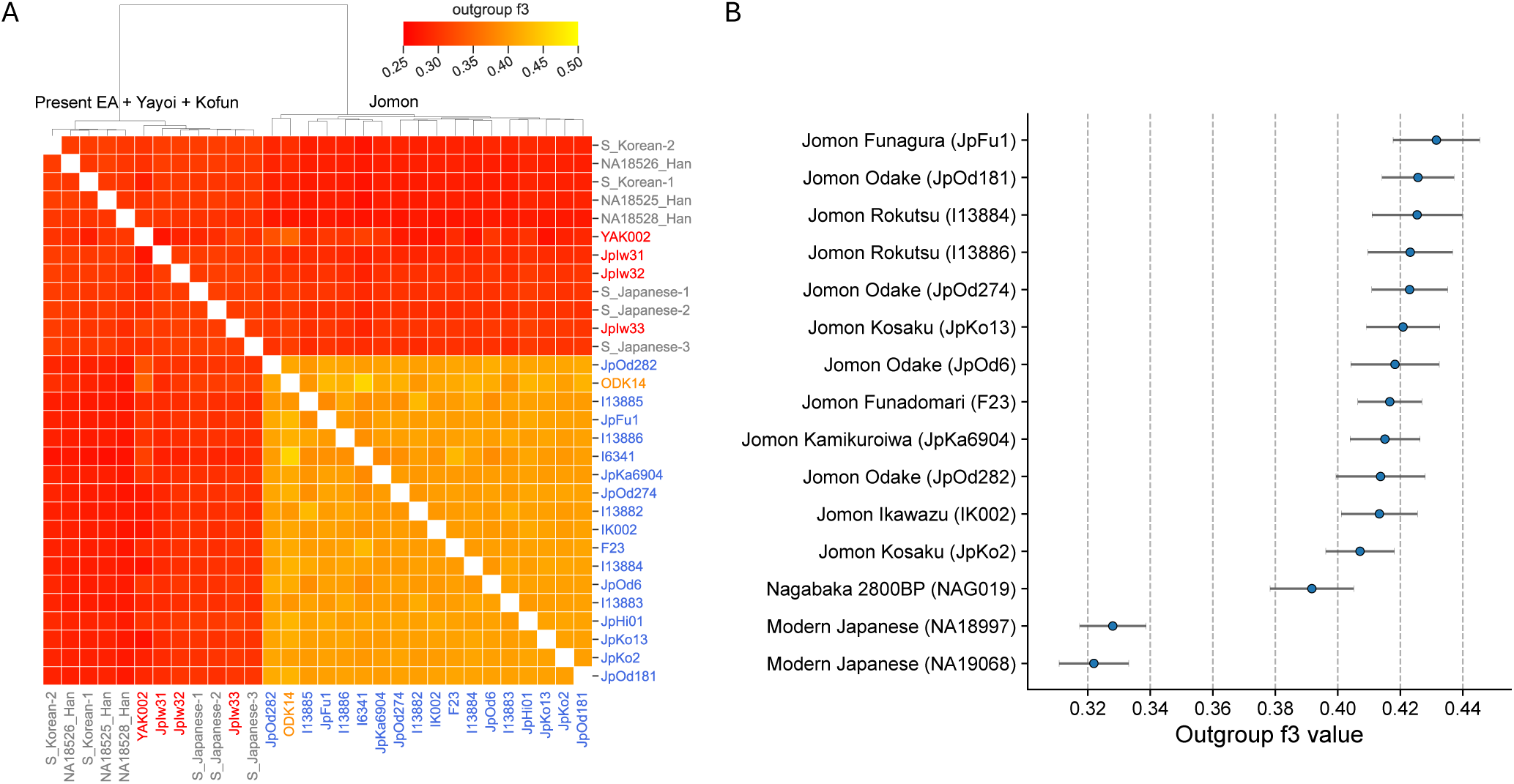
Results of outgroup-*f3* statistics for ODK14. This figure summarizes outgroup-*f3* values for the ODK14 individual using Mbuti as the outgroup, in comparison with ancient and present-day East Asian populations from the Japanese archipelago and neighboring regions. (A) Hierarchical clustering based on outgroup-*f3* values. The color bar represents the magnitude of *f3* values, and the dendrogram illustrates clustering of individuals with similar pairwise *f3* values. (B) The 15 individuals with the highest outgroup-*f3* values among those sharing at least 5,000 SNPs with ODK14 in the population panel dataset. Error bars indicate standard errors.

**Figure 5.**
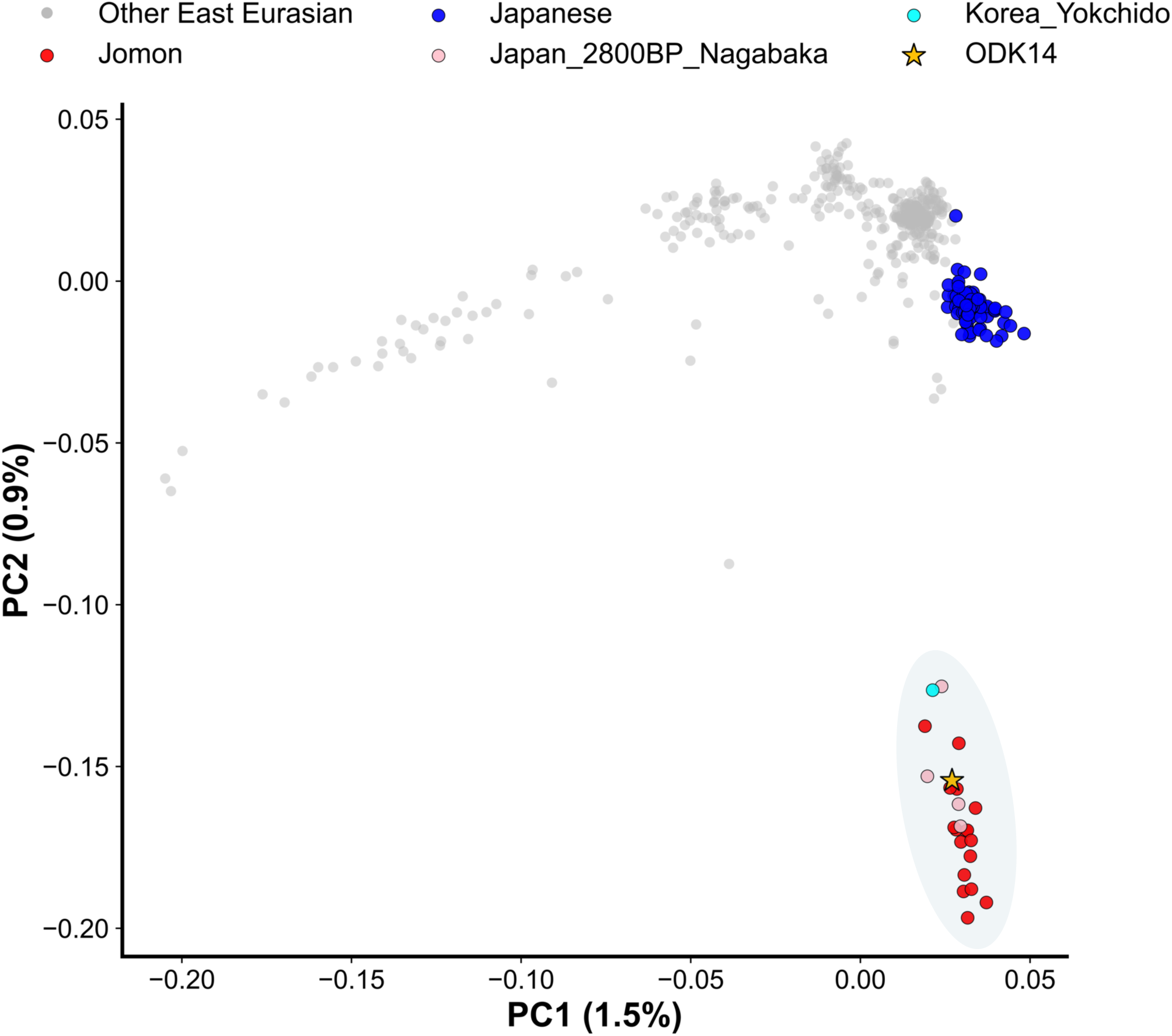
Pricipal component analysis with East Eurasian populations. This figure illustrates the distribution of East Eurasian populations along the first two principal components (PC1 and PC2) derived from PCA using a reference panel of East Eurasian groups. The percentages on each axis represent the proportion of genetic variance explained by the corresponding component. Individuals from the Japanese Archipelago, both present-day and ancient, are highlighted in color, while other East Eurasian populations are shown in gray for comparison. Colored shaded areas indicate clusters that include individuals classified as Jomon cluster based on outgroup-*f3* statistics.

**Figure 6.**
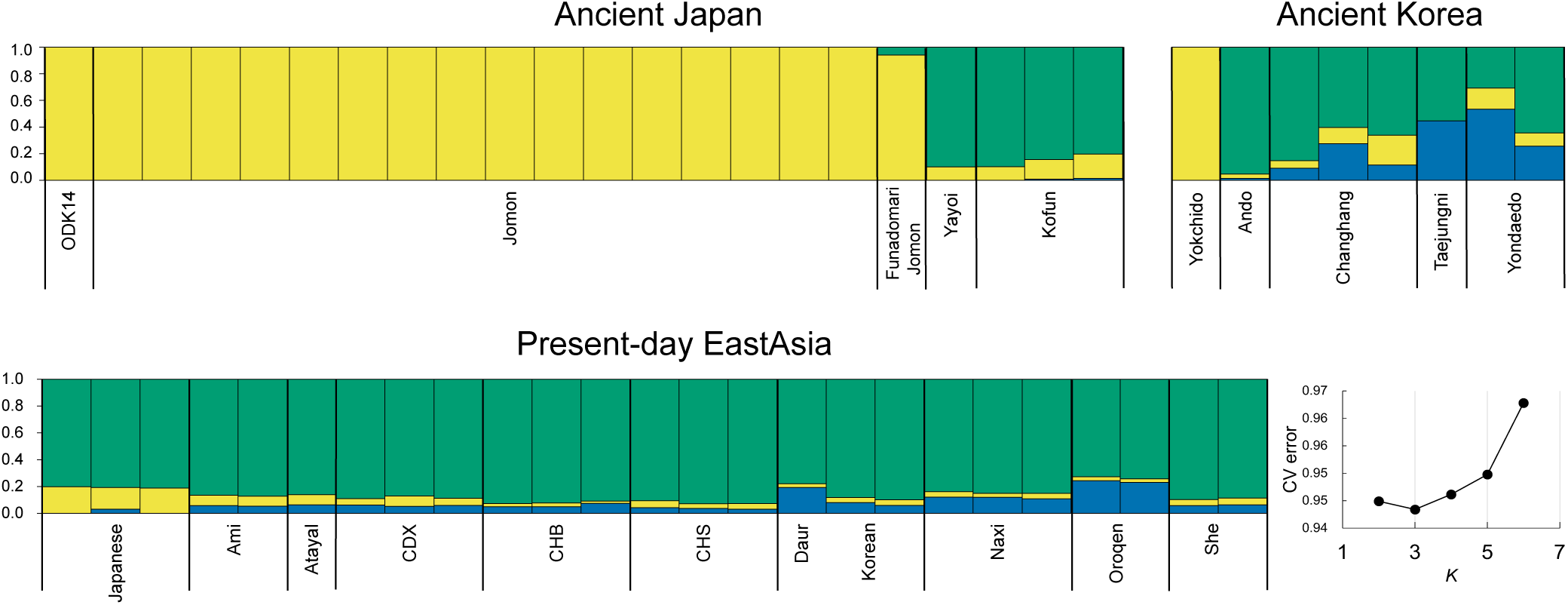
Selected individuals from a ADMIXTURE analysis. The bar plots display individuals from the Japanese Archipelago and surrounding regional populations based on ADMIXTURE analysis using a reference panel dataset. Ancestry components were estimated for values from *K*=2 to *K*=10 with 10-fold cross-validation; as shown in the bottom-right inset, *K*=3 yielded the lowest cross-validation error. Accordingly, the bar plots displays the ADMIXTURE results at *K*=3, where components P1, P2, and P3 are represented in green, yellow, and blue, respectively. The results at *K*=3 for all individuals are provided in Figure S7.

Together, these analyses indicate that ODK14—sequenced using ONT technology—shows a population genetic profile that aligns closely with those of previously reported Jomon individuals using Illumina sequencing [47–49].

## 3. Discussion

This study presents the first successful application of Oxford Nanopore Technologies (ONT) to ancient DNA (aDNA) analysis, demonstrating that nanopore sequencing can recover authentic aDNA signatures and generate population genetic results highly concordant with those obtained from Illumina data. Despite being produced through fundamentally different sequencing chemistries, ONT-derived reads preserved key molecular indicators of aDNA—such as terminal deamination and extensive fragmentation—and showed low levels of exogenous contamination. Furthermore, mitochondrial haplogroup assignment, genome-wide depth estimates, and transversion-site genotypes displayed high concordance between ONT and Illumina platforms (Table 1), and population genetic analyses placed ODK14 alongside previously reported Jomon individuals (Fig. 4-6). These findings collectively demonstrate that ONT sequencing can produce reliable aDNA datasets suitable for downstream genomic analyses.

Beyond its compatibility with aDNA, ONT sequencing offers several unique advantages for paleogenomic applications. ONT platforms enable long-read sequencing, real-time data acquisition, and fully portable operation, making them particularly well suited for use outside traditional laboratory environments. Our time-resolved analysis showed that informative genomic signals—such as alignment yield (Fig. 1B), SNP recovery (Fig. 1B) and sex inference (Fig. 3)—emerged rapidly during sequencing, with sex determination stabilizing within the first hour. Such rapid data acquisition highlights the potential for ONT to support on-site paleogenomics, including preliminary assessment of archaeological samples, triaging of specimens for deeper sequencing, and rapid decision-making in remote or logistically constrained field settings.

However, several technical limitations remain in aDNA sequencing using ONT platforms. Differences in damage-repair treatments must be considered, yet ONT-derived variants did not achieve complete concordance with Illumina data even when restricted to transversion sites (Table 1). In addition, we observed novel singletons within the same mitochondrial haplogroups (Fig. S5). These discrepancies may reflect the inherent challenges of aligning ultra-short and damaged fragments using nanopore sequencing technologies originally optimized for high-molecular-weight DNA, as well as sequencing and basecalling characteristics associated with the platform.

The number of usable reads was also constrained by the active pore count and showed pronounced output decay during extended sequencing runs (Fig. 1A). In our dataset, approximately 90% of total reads were produced in the first 48 hours, with markedly reduced output thereafter; thus, continuous 72-hour runs may not be optimal. Furthermore, approximately 67% of reads exhibited translocation speeds slower than expected (Fig. S2), likely due to residual constrictive debris remaining after purification, DNA damage (including fragmentation and physical denaturation), adapter ligation artifacts, or partial pore blockage, all of which may contribute to delayed translocation through the nanopores and, consequently, a reduction in overall sequencing throughput (Fig. 1A). Therefore, incorporating flow cell wash procedures or segmentation of sequencing sessions could help sustain throughput.

A key distinction between ONT and short-read platforms such as Illumina lies in the intrinsic calculation of per-base quality scores, which may affect downstream variant calling. In ancient DNA analyses, post-hoc damage-aware trimming (masking) or quality rescaling can help mitigate these effects, while continued improvements in basecalling models and signal-level damage-aware algorithms are expected to further enhance accuracy. Emerging laboratory strategies—such as ultra-short DNA library preparations [50], concatenation-based fragment reconstruction [51], and optimized adapter chemistries—are likely to increase ONT’s suitability for analyzing extremely degraded materials. Adaptive sampling technologies [52, 53] may one day permit real-time enrichment of aDNA, though current fragment-length constraints remain a barrier.

Despite these limitations, the broader implications of this study are substantial. The portability, minimal infrastructure requirements, and real-time data output of ONT platforms create a pathway toward on-site paleogenomics, enabling sequencing at archaeological excavations, regional museums, and local research institutions. This shift has far-reaching consequences for global scientific equity. In many regions, the export of archaeological remains involves extended administrative procedures that can take several months or even years, while in some areas it is subject to strict limitations or may not be permitted. Unauthorized export may result in significant repercussions, potentially limiting opportunities for international genomic collaboration. ONT-based sequencing circumvents these challenges by allowing local researchers to generate genomic data without relinquishing physical control of skeletal materials, preserving cultural heritage sovereignty and enabling countries to retain scientific ownership of their ancestral remains.

By doing so, nanopore-based onsite sequencing offers a feasible approach to addressing long-standing concerns among archaeologists and geneticists regarding sample accessibility, authorship, and research leadership. By empowering local archaeologists and anthropologists to independently generate and analyze aDNA data, ONT sequencing may accelerate the global accumulation of ancient genomes, encourage more equitable scientific collaboration, and reshape ethical norms in paleogenomics.

In summary, while further optimization of wet-lab protocols, sequencing chemistries, and computational frameworks is needed, our results establish ONT sequencing as a technically robust, operationally flexible, and ethically transformative tool for ancient DNA research. Nanopore platforms have the potential to usher in a new era of rapid, field-deployable, and socially sustainable paleogenomics, enabling genomic analysis of archaeological human remains in ways that were previously inaccessible.

## 4. Materials and Methods

### 4.1 Archaeological sample

The skeletal remains analyzed in this study derive from individual ODK14, excavated from the Odake shell mound in Toyama Prefecture, located along the northern coast of central Honshu, Japan. The site is attributed to the Early Jomon period based on archaeological stratigraphy and radiocarbon dating. A dense region of the petrous portion of the temporal bone was selected for ancient DNA extraction due to its high endogenous DNA preservation properties. All destructive sampling procedures were conducted under institutional approval and in accordance with ethical guidelines for the study of archaeological human remains.

### 4.2 Experimental workflow

All laboratory procedures were carried out in a dedicated ancient DNA clean-room facility that is physically isolated from post-PCR laboratories and equipped with positive air pressure, HEPA filtration, and UV decontamination systems. Personnel wore full-body lab wears, face shields, and gloves, and all work surfaces and instruments were treated with diluted sodium hypochlorite, DNA decontaminants (DNA-OFF (TaKaRa) and DNA AWAY (Thermo Fisher Scientific)) and UV irradiation to minimize modern DNA contamination. For sampling, approximately 50 mg of bone powder was obtained from the inner surface layer of cochlea at the clean bench using a sterilized dental diamond disc and tungsten carbide bur (Shofu) operated at low rotational speed. The powder was collected into 2.0 ml DNA Lobind tube (Eppendorf) and transferred immediately to the extraction workflow.

DNA extraction followed a silica-based protocol with D buffer optimized for highly fragmented ancient DNA [54]. DNA extraction was carried out using Biomeck i5 (Beckman Coulter), the automation robot in Kanazawa University. For the ONT dataset, DNA was extracted without uracil-DNA glycosylase (UDG) treatment in order to preserve deamination signatures needed for damage assessment. For the Illumina dataset, a separate aliquot underwent UDG treatment to remove deaminated cytosines and thereby reduce the frequency of C > T substitution, postmortem damage on variant calling. DNA concentrations were measured using Qubit dsDNA High Sensitivity assays (Thermo Fisher Scientific), and fragment-length distributions were evaluated using High Sensitivity D1000 Kit of Agilent 4200 TapeStation System (Agilent) to confirm the expected short-fragment profile of ancient DNA.

Library preparation for ONT sequencing was performed using NEBNext Ultra II DNA Library Prep Kit for Illumina (New England Biolabs), NEBNext Multiplex Oligos for Illumina (96 Unique Dual Index Primer Pairs) (New England Biolabs), Ligation Kit (SQK-LSK114) (Oxford Nanopore Technologies) and NEBNext Companion Module v2 for Oxford Nanopore Technologies Ligation Sequencing (New England Biolabs). First, NEB library preparation with beads size selection was applied so as to exclude long DNA fragments, mondern DNA contaminants [48]. Second, ONT library preparation was conducted with Ligation Kit and Short Flagment Buffer to attach the DNA adapter with morter protein. For Illumina sequencing, four double-stranded libraries (ODK14_UDG_1–4) were constructed using the NEBNext Ultra II DNA Library Prep Kit for Illumina (New England Biolabs), following established ancient DNA protocols that include USER treatment [48]. Libraries were purified using SPRI beads solution (Beckman Coulter) and quantified using Qubit and Tapastation analysis before sequencing. The UDG-treated Illumina dataset was used as a validation dataset for cross-platform comparison with the ONT-derived data.

### 4.3 Sequencing

We used the PromethION 2 Solo (P2 Solo; Oxford Nanopore Technologies) equipped with R10.4.1 flow cells to perform DNA sequencing. We assessed flow cell quality ensuring more than 5,000 active pores and obtained raw squiggle data (POD5 format) with MinKNOW (MKE_1013_v1_revDK_25Apr2025). Then, we basecalled the squiggle data using Dorado (v1.0.2) (https://github.com/nanoporetech/dorado) with the super accurate model (dna_r10.4.1_e8.2_400bps_sup@v5.2.0) and generated BAM files. Dorado also detected DNA modifications, most notably 5-methylcytosine (5mC), from the raw nanopore signal. Finally, we converted the resulting BAM files to FASTQ format using the SamToFastq command in Picard tools (v3.4.0) (http://broadinstitute.github.io/picard/) and used these FASTQ files for downstream analyses. We also prepared four Illumina NGS libraries (ODK14_UDG_1 to ODK14_UDG_4) using the NEBNext Ultra II DNA Library Prep Kit and sequenced them on the NovoSeq 6000 platform with 151-cycle paired-end reads and 8 bp dual index sequences in Kyoto University.

### 4.3 Data processing for sequencing data

We first evaluated ONT sequencing reads and the corresponding sequencing summary using Sequali (v1.0.1) [55]. We then filtered low-quality ONT reads with NanoFilt (v2.8.0) [56], removing sequences with an average Phred score below 15 or a length shorter than 30 bp. Adapter trimming was performed with Porechop (v0.2.4) (https://github.com/rrwick/Porechop/), and reads shorter than 30 bp after trimming were discarded using SeqKit (v2.10.0) [57]. The remaining reads were used for downstream genome alignment. For Illumina NovoSeq 6000 paired-end data, we trimmed adapters and low-quality bases and merged overlapping pairs (minimum overlap: 11 bp) using AdapterRemoval (v2.3.4) [58], followed by removal of reads shorter than 30 bp prior to alignment.

We aligned quality-controlled reads from both sequencing platforms to the human reference genome (hg19/GRCh37) using BWA aln (v0.7.18) with parameters optimized for ancient DNA (-l 1024 -n 0.01 -o 2). We removed duplicate reads with Picard MarkDuplicates (v3.4.0) and discarded alignments with mapping quality <30 using Samtools (v1.16.1) [59].

To classify ONT reads by sequencing runtime, we extracted read identifiers and timestamps from Dorado summary files (v1.0.2). We grouped reads into 1-minute intervals during the initial 0–100 minutes, into 1-hour intervals from 1–10 hours, and into 10-hour intervals from 10–80 hours. Because the sequencing runs lasted approximately 72 hours, the 80-hour upper boundary encompassed all reads generated during the run.

To authenticate aDNA in ONT sequencing data, we analyzed UDG-untreated ONT alignments after duplicate removal. We quantified terminal deamination and fragmentation patterns with mapDamage (v2.2.3) [60] and examined mismatch edit-distance profiles with damageProfiler (v1.1) [61]. We estimated contamination levels using MitoSuite (v1.0.9) [62], based on mitochondrial variation, and ANGSD (v0.940) [63], based on X-chromosomal polymorphisms.

To reduce the impact of damaged-derived substitutions during variant calling, we applied two preprocessing steps to ONT reads. First, we rescaled base quality scores at damage-prone positions using the --rescale option in mapDamage (v2.2.3) [60], which allowed subsequent quality filtering to downweight or remove deaminated sites. Second, for libraries exhibiting substantial terminal damage, we trimmed 2 bp from both ends of each read using bamUtil (v1.0.15) [64]. We used these damage-mitigated ONT alignments for downstream genotyping. Finally, we obtained genotypes for the 1240K SNP panel using pileupcaller (sequenceTools v1.6.0.0) (https://github.com/stschiff/sequenceTools) in random haploid mode.

### 4.4 Biolocal sex inference

To infer the biological sex of the sample ODK14, we used the Ry [43] defined as:

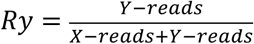

where Y-reads and X-reads represent the number of reads aligned to the human Y and X chromosomes, respectively. We also tested whether time-course sex determination is feasible during ONT sequencing by using only reads generated during the initial 0–100 minutes of the run.

### 4.4 Merging to published datasets

We merged the newly generated genotype data from individual ODK14 with published datasets using PLINK (v1.90b6.21) [65]. We selected comparative present-day and ancient individuals primarily from East Eurasia based on the 1240K SNP panel of the Allen Ancient DNA Resource (AADR v54.1) [66]. We further incorporated recently reported ancient individuals from East Asia [67] into this merged dataset. To minimize the impact of ancient DNA damage on downstream population-genetic analyses, we restricted all analyses to transversion sites. We also estimated pairwise relatedness using READ (v2.1.1) [68] and excluded individuals identified as first-degree relatives within any population group. The final designed panel dataset is summarized in Supplementary Table 1.

### 4.4 Maternal/Paternal lineage inference

To estimate the mtDNA lineage of ODK14, we inferred the maternal mitochondrial haplogroup from QC-filtered alignment reads using MitoSuite (v1.0.9) [62] and generated a full-length mitochondrial consensus sequence. We used this consensus sequence, together with a comparative dataset comprising present-day Japanese, published Jomon individuals, and related haplogroup genomes (Table S1), to construct a median-joining network in PopART (v1.7) [69]. We also analyzed Y-chromosomal variation using Y-LineageTracker (v1.3.0) [70] for estimating paternal lineage of ODK14.

### 4.5 Principal component analysis

To investigate the genetic relationship between the Jomon individual ODK14 and neighboring populations, we used the merged comparative panel dataset (Table S2), which includes populations and individuals primarily from East Eurasia. Before performing PCA, we reduced redundancy among SNPs due to linkage disequilibrium (LD) by conducting LD pruning in PLINK (v1.90b6.21) [65] with the parameters “--indep-pairwise 50 10 0.1”. We then ran smartPCA in EIGENSOFT (v8.0.0) [45] on the LD-pruned dataset to infer population structure.

### 4.6 Outgroup-f3

To assess genetic affinities between pairs of individuals (X1 and X2), we obtained outgroup-*f3* statistics from the panel dataset using ADMIXTOOLS (v7.0.2) [44]. We calculated *f3*(X1, X2; Mbuti), with Mbuti as the outgroup, to quantify the degree of shared genetic drift between each pair of individuals. Based on these results, we identified the individuals showing strong affinity to ODK14. To further visualize relationships among individuals, we performed hierarchical clustering—based on Euclidean distances derived from their pairwise outgroup-*f3* values—using a dataset consisting of previously reported Jomon individuals, post-Yayoi individuals, and present-day East Asians.

### 4.7 Admixture

To characterize genome-wide ancestry patterns in ODK14, we applied an unsupervised model-based clustering approach implemented in ADMIXTURE (v1.3.0) [46]. Before running the analysis, we performed LD pruning to reduce the influence of linked SNPs, using the same filtering parameters as in the PCA dataset. We explored *K* values from 2 to 10 and carried out 10-fold cross-validation for each *K* to identify models that best captured population substructure. Independent runs showed consistent convergence, and we selected the solutions with the lowest cross-validation errors. We then compared the resulting ancestry profiles with those of previously published Jomon individuals to evaluate whether ODK14 carries ancestry components characteristic of Jomon-related populations.

## Acknowledgements

This work was supported by JSPS KAKENHI Grant Number 19K16246, 20H05822, 22H02711, Kanazawa University SAKIGAKE Project, and MEXT Program for Forming Japan’s Peak Research Universities (J-PEAKS). The super-computing resource was provided by Human Genome Center (the Univ. of Tokyo). We thank Single-cell Genome Information Analysis Core (SignAC) at WPI-ASHBi, Kyoto University for their support.

## Supplementary Files

**Fig. S1.**
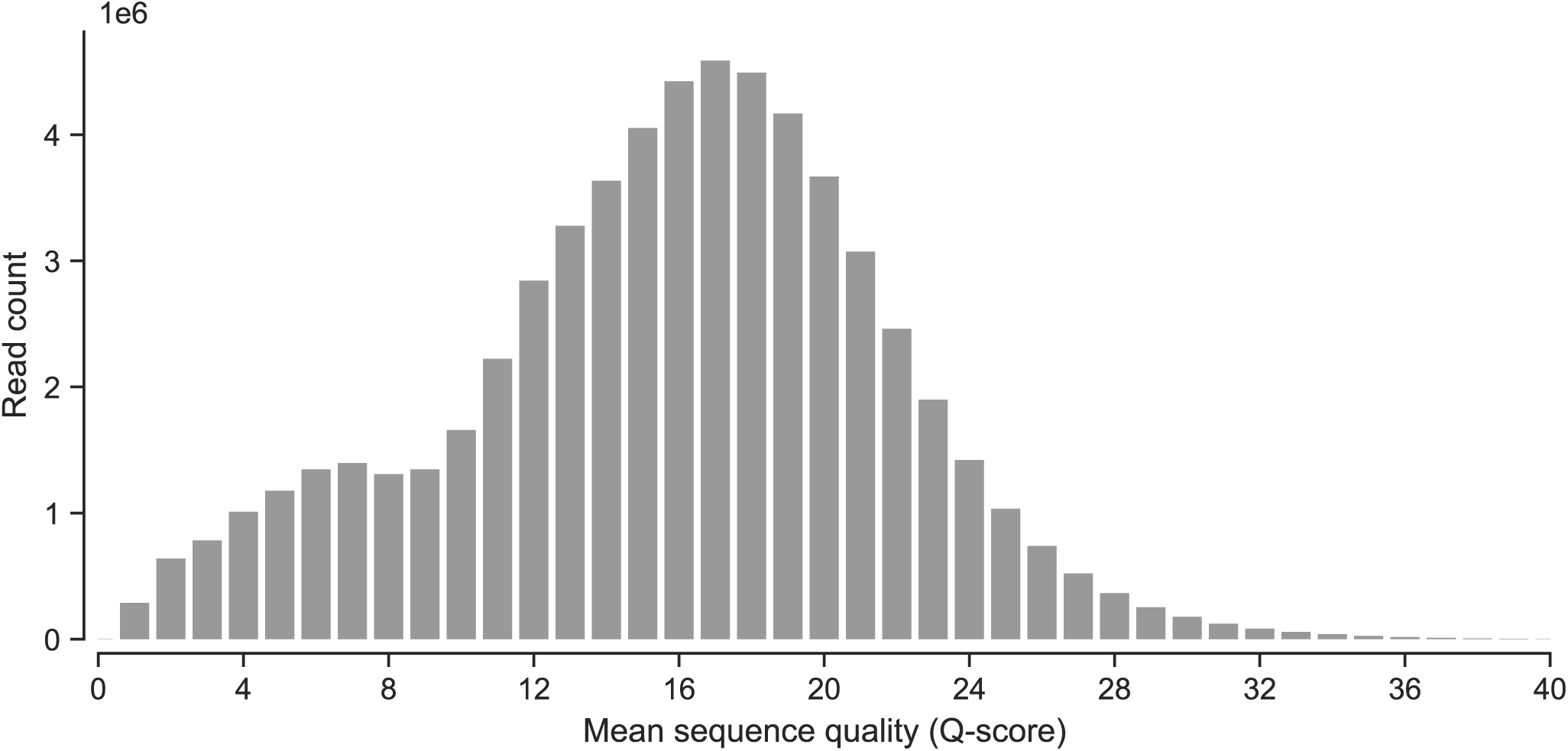
Distribution of per raw-sequence quality scores from ONT sequencing data.

**Fig. S2.**
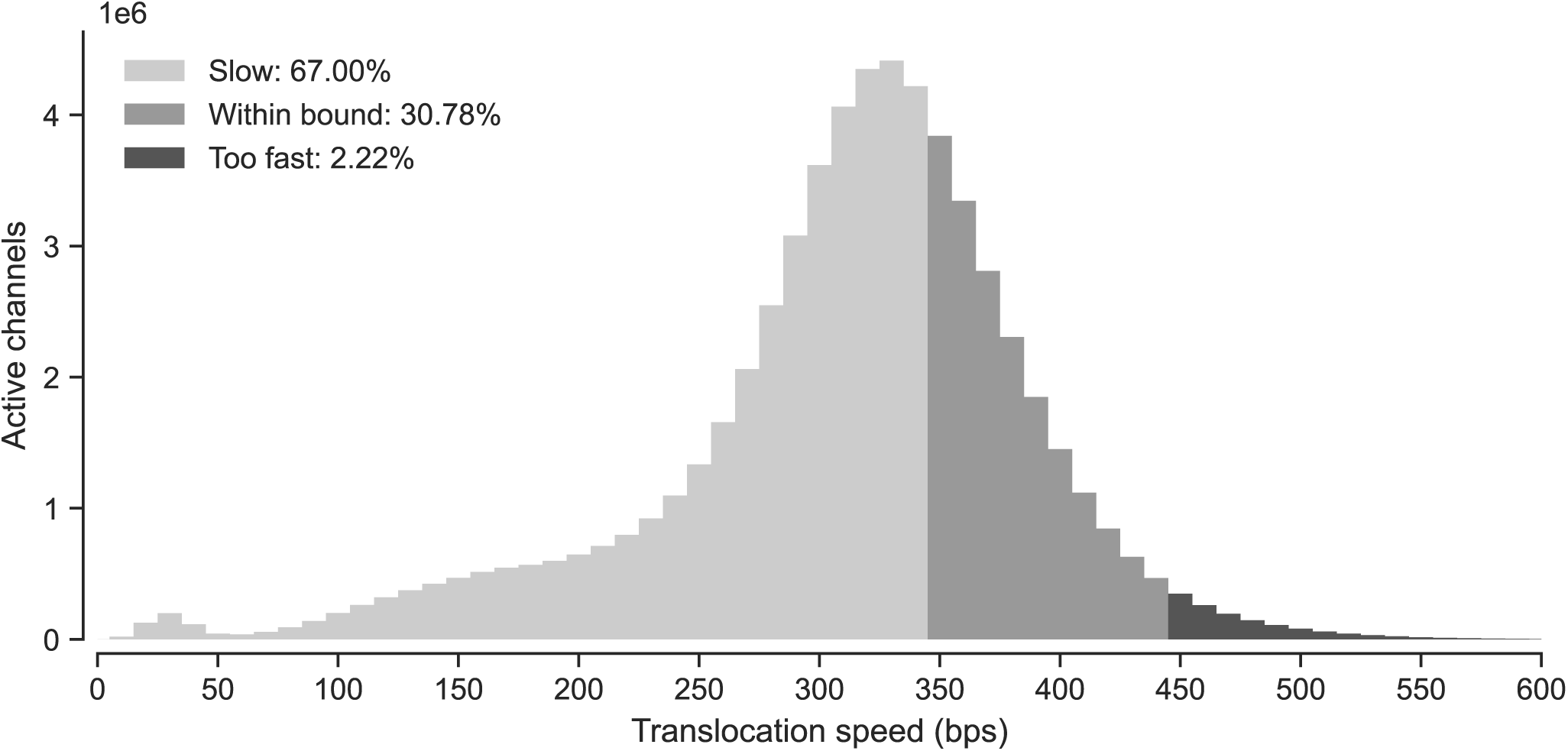
Distribution of nanopore translocation speeds across active channels.

**Fig. S3.**
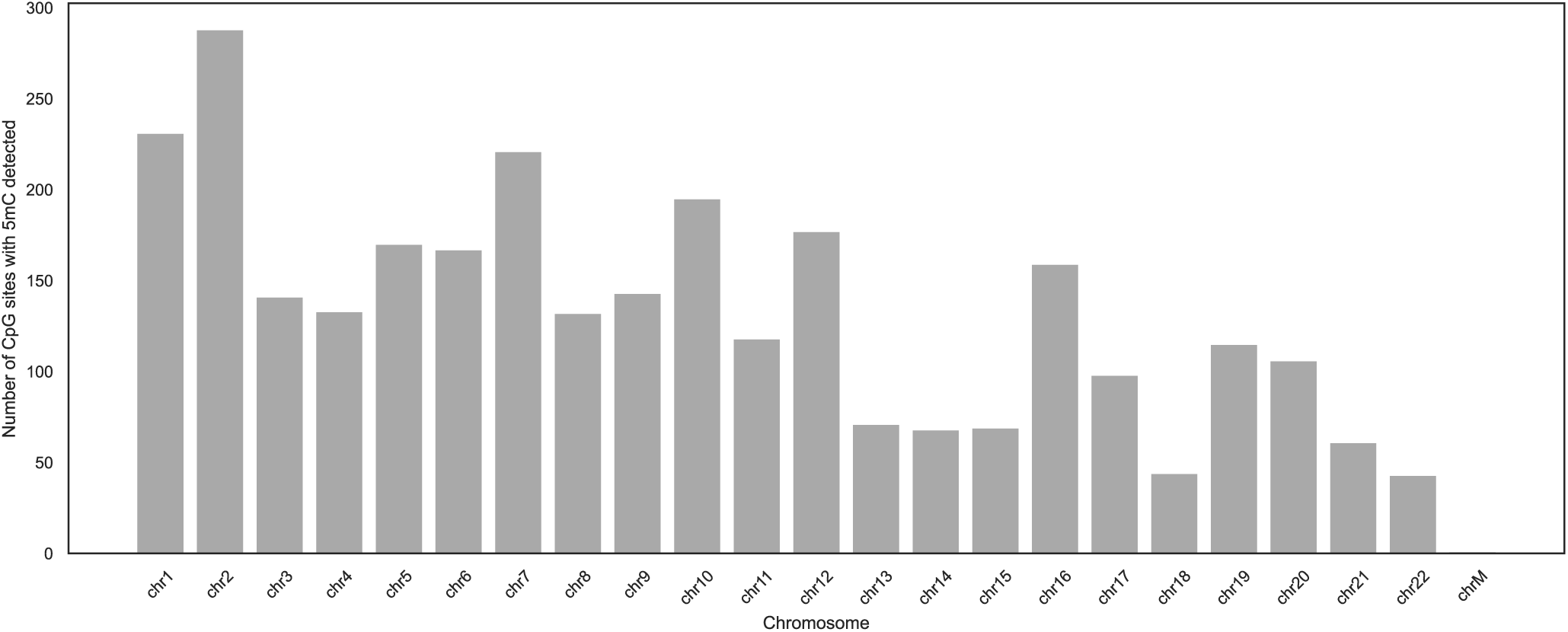
Detected 5mC at CpG sites across each chromosome.

**Fig. S4.**
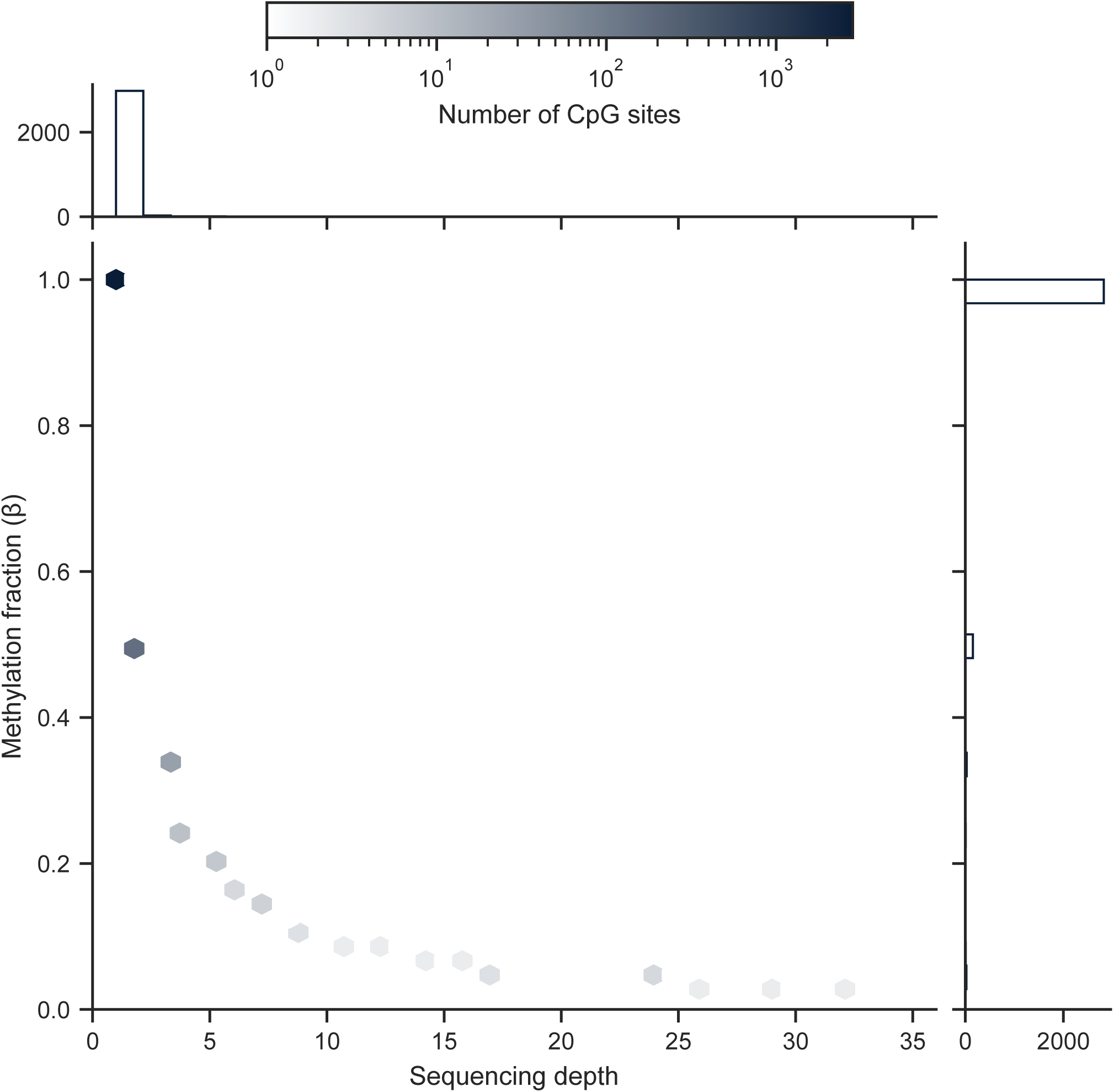
Joint distribution of sequencing depth and CpG methylation fraction.

**Fig. S5.**
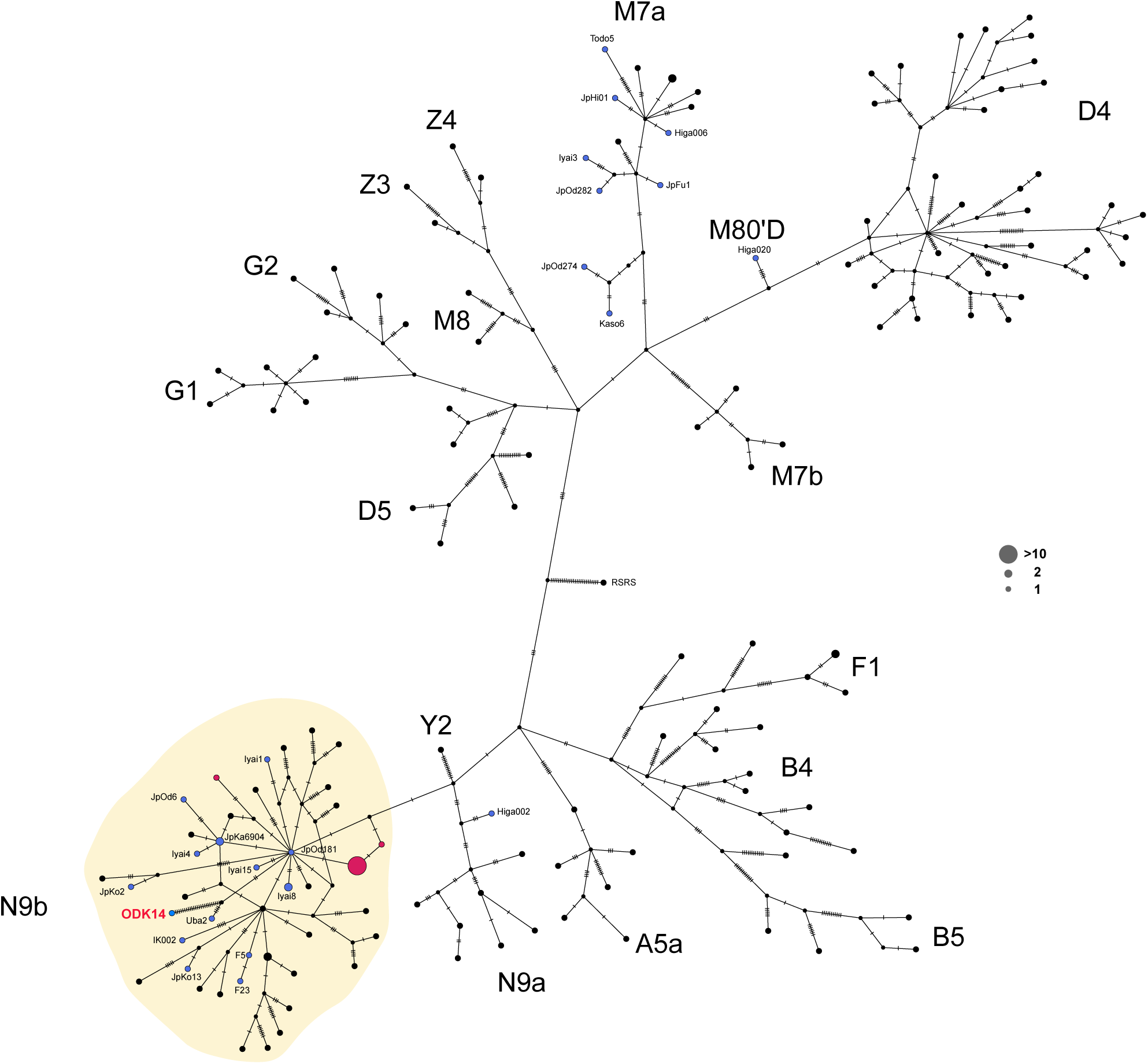
Median-joining network of mitochondrial DNA from present-day Japanese, Siberian individuals, and ancient Jomon samples. This network was constructed using mtDNA sequences from modern Japanese, modern Siberian individuals from Russia, and ancient Jomon samples. Circles represent individual sequences or groups of individuals sharing identical haplotypes. Circle colors correspond to population categories: black for present-day Japanese, magenta for Siberians, and blue for Jomon individuals. Circle size reflects the number of individuals within each haplotype. Hatch marks along the connecting branches indicate the number of single mutational steps between haplotypes. Bold labels denote mitochondrial haplogroups or sub-haplogroups. Shaded regions indicate nodes belonging to haplogroup N9b.

**Fig. S6.**
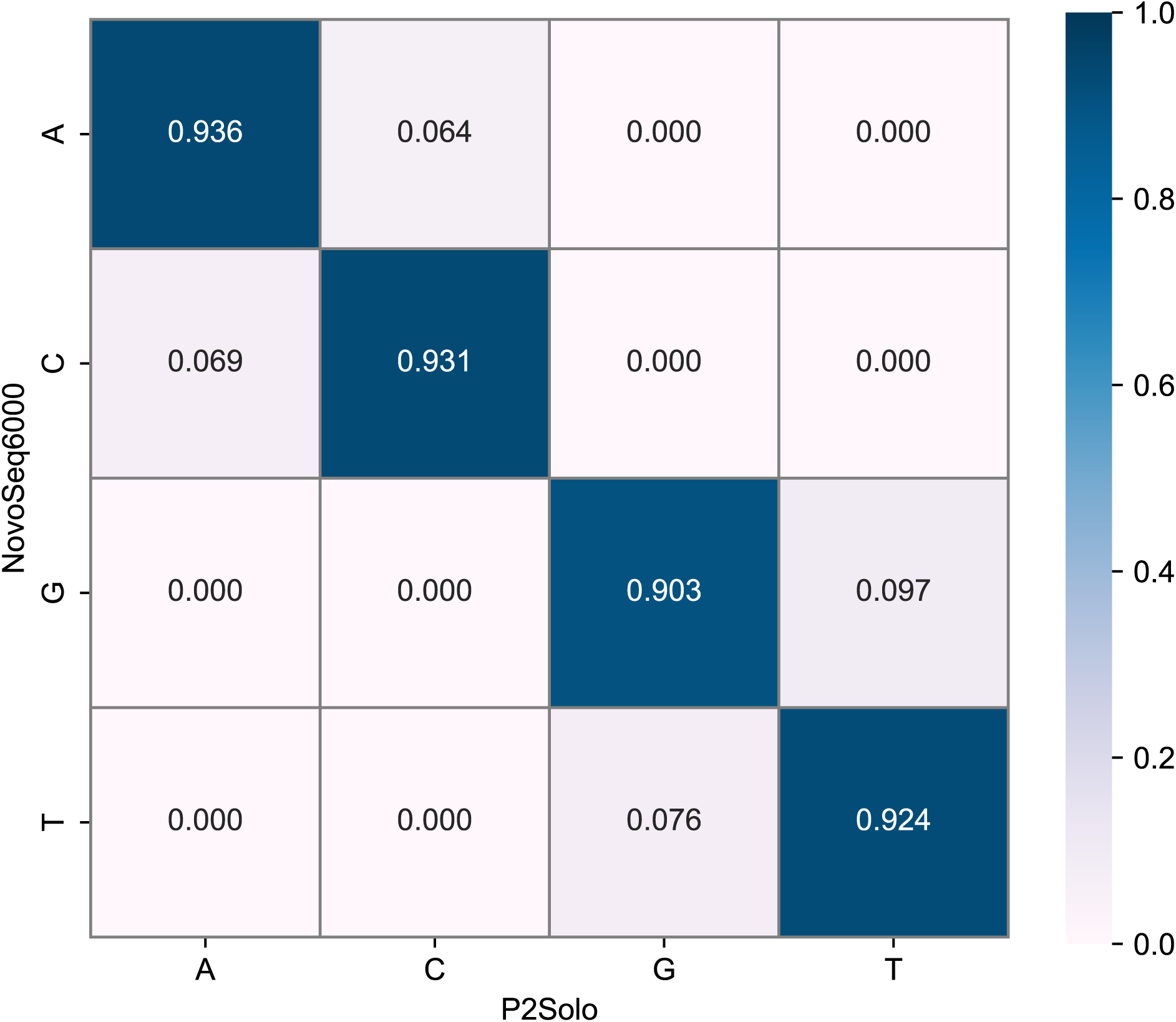
Variant concordance between NovoSeq6000 and P2Solo at transversion sites.

**Fig. S7.**
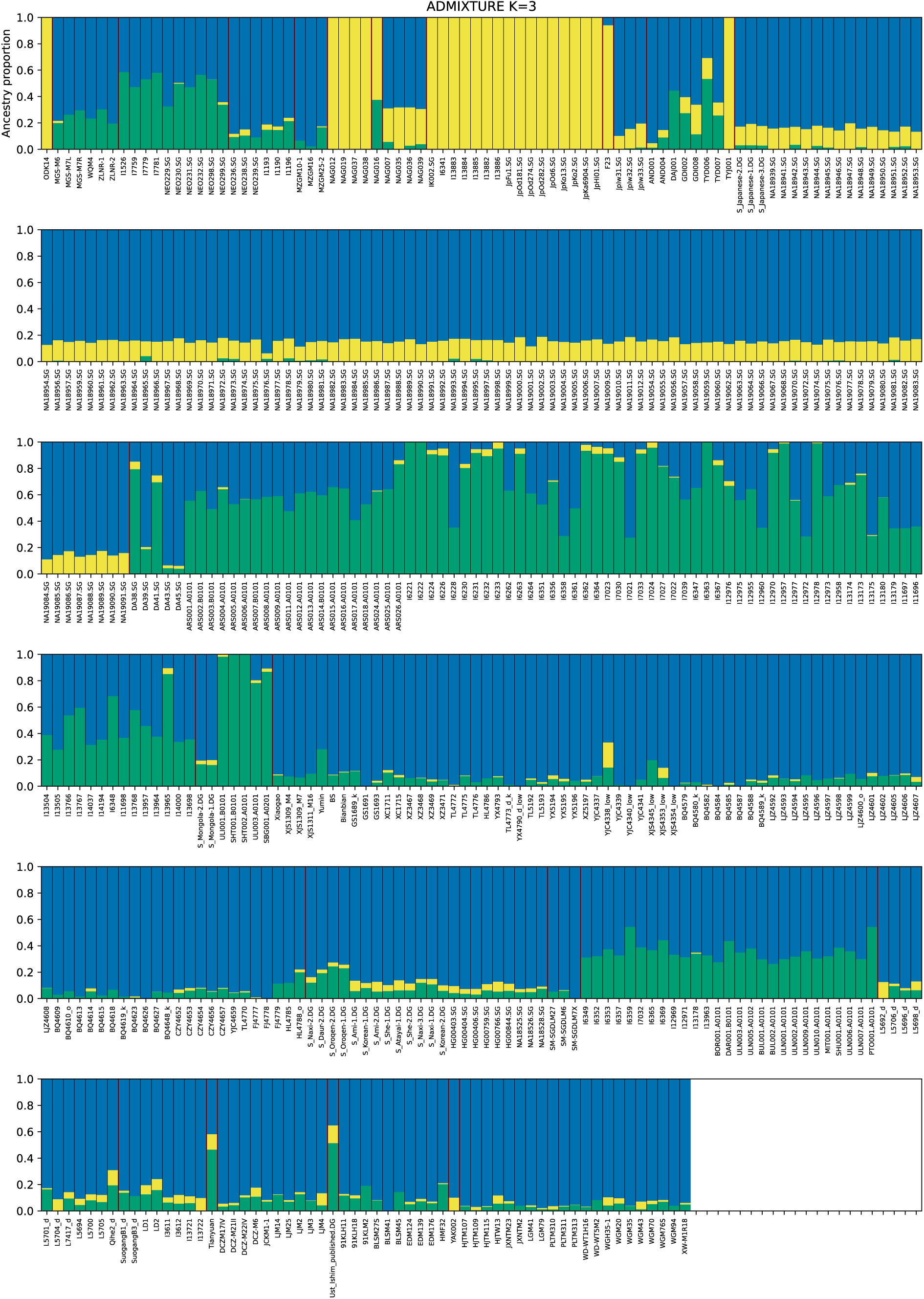
Results of the ADMIXTURE analysis (K=3).

